# Functional characterization and molecular engineering of a *O*-methyltransferase involved in bis-benzylisoquinoline alkaloids biosynthesis from *Nelumbo nucifera*

**DOI:** 10.64898/2026.02.05.703910

**Authors:** Yuetong Yu, Xinyi Qi, Hedi Zhao, Zhennan Wang, Sha Chen

## Abstract

*Nelumbo nucifera* (lotus), a traditional aquatic plant in Asia, is valued for its nutritional and therapeutic properties. Benzylisoquinoline alkaloids (BIAs) are its major bioactive components, with significant bioactivities and pharmacological values. O-methyltransferases (OMTs) play a crucial role in shaping the structural and functional diversity of BIAs. Herein, we characterized a specific OMT, designated Nn7OMT, which exhibits a novel catalytic function: methylating the C7 position of the bisbenzylisoquinoline (bisBIA) skeleton. Nn7OMT also displayed substrate promiscuity, catalyzing O-methylation at the C7 position of bisBIAs, C6/C7 positions of 1-benzylisoquinolines, C7 position of aporphines, and C9 position of protoberberines, with isoliensinine as its preferred substrate. The expression profile of Nn7OMT correlates with bisBIA accumulation in planta, supporting its involvement in bisBIA biosynthesis in *N. nucifera*. Protein engineering guided by molecular docking and molecular dynamics simulations identified key residues critical for Nn7OMT activity. Mutants M158A and V306A retained catalytic activity toward isoliensinine while showing nearly undetectable activity against 1-benzylisoquinoline alkaloids, thereby significantly improving Nn7OMT’s substrate specificity. These findings advance our understanding of BIAs biosynthesis in lotus and provide valuable biocatalysts with enhanced specificity.

**SIGNIFICANCE:** In this study, we report the innovative discovery and characterization of Nn7OMT, the first O-methyltransferase isolated from Nelumbo nucifera that harbors a novel catalytic activity toward the bisbenzylisoquinoline alkaloid (bisBIA) backbone. Nn7OMT displayed substrate promiscuity, catalyzing methylation at the C7 position of the bisbenzylisoquinoline skeleton (isoliensinine), C6/C7 positions of the 1-benzylisoquinoline skeleton (norcoclaurine, coclaurine, N-methylcoclaurine), C7 position of the aporphine skeleton (lirinidine), and C9 position of the protoberberine skeleton (scoulerine)—with isoliensinine as its most preferred substrate. Innovatively, the catalysis of isoliensinine by Nn7OMT was a novel function and the first O-methyltransferase identified to catalyze the bisbenzylisoquinoline backbone, which may be involved in the biosynthesis of neferine. Nn7OMT’s expression profile correlated with bisBIA accumulation in planta, further supporting its role in bisBIA biosynthesis in *N. nucifera*. We further engineered Nn7OMT using molecular docking and MD simulation-guided strategies. Mutants M158A and V306A retained catalytic activity for isoliensinine while showing nearly undetectable activity against 1-benzylisoquinoline alkaloid substrates. This targeted engineering significantly improved the specificity of Nn7OMT, rendering it highly suitable for methylated isoliensinine biosynthesis by reducing by-product accumulation and simplifying purification processes. Our work deepens understanding of the unique BIA metabolic pathway in *N. nucifera* through the identification of the first bisBIA-catalyzing O-methyltransferase, and provides valuable biocatalysts with enhanced substrate specificity.

## INTRODUCTION

*Nelumbo nucifera* (lotus) is an important dual-purpose crop for medicine and food. It has been widely used in Asia for more than 2000 years, owing to its high nutritional, ornamental, and medicinal values ^1^. Different parts of the lotus, including leaves, plumules, seeds, and roots, exhibit various biological and pharmaceutical activities ^2^. Traditionally, these parts are used to treat obesity, diarrhea, high fever, insomnia, hypertension, arrhythmia, hyperlipidemia, and neurological disorders^3,4^. Benzylisoquinoline alkaloids (BIAs) are one of its primary bioactive components, mainly including three structural skeletons of monobenzylisoquinolines, aporphines, and bisbenzylisoquinolines^5^. BIAs in lotus possessed important bioactivities and pharmacological values. For example, the two monobenzylisoquinoline alkaloids norcoclaurine and coclaurine exhibit anti-HIV activity ^6^. Armepavine is a potential therapeutic treatment for autoimmune illnesse such as autoimmune crescentic glomerulonephritis and systemic lupus erythematosus ^7,8^. The aporphine alkaloid nuciferine exhibits antidiabetic, anti-HIV, anti-melanogenesis, and anticancer properties ^9,10^. Neferine, a bisbenzylisoquinoline alkaloid abundant in lotus plumules, has been reported to have therapeutic potential against COVID-19 ^11^. It also exerts multiple therapeutic effects, including anti-inflammatory, antioxidant, antihypertensive, anti-arrhythmic, antiplatelet, antithrombotic, anti-amnesic, and negative inotropic effects, as well as antitumor activities against lung, liver, and breast cancers ^12,13^.

BIAs metabolism has been intensively investigated in Ranunculales species^14^. Most of their biosynthetic pathways have been fully elucidated, such as the pathways responsible for the synthesis of morphine, berberine, and sanguinarine^15-17^. Notably, all BIAs are derived from L-tyrosine and proceed via (S)-norcoclaurine, the first intermediate and a common precursor for all BIAs ^18^. While distinct structural subclasses have evolved and are distributed across different plant species. In *N. nucifera*, BIA metabolism initiates with norcoclaurine formation, followed by the generation of structurally diverse 1-benzylisoquinolines via a set of enzymes, such as O-methyltransferases (OMTs) and N-methyltransferases. Subsequently, 1-benzylisoquinolines are converted into aporphines and bisbenzylisoquinolines via intramolecular C-C phenol coupling and intermolecular C-O phenol coupling, which are catalyzed by the CYP80G and CYP80A subfamily enzymes, respectively ^19^. Currently, a study elucidated the biosynthetic pathway of pronuciferine and bis-benzylisoquinoline catalyzed by NnCYP80Q1 and NnCYP80Q2, respectively, in *N. nucifera* ^20^. Two *O*-methyltransferases from *N. nucifera* have been identified which catalyze the 6-*O*- and 7-*O*-methylation of the 1-benzylisoquinoline backbone ^21^. Our recent study further revealed that an O-methyltransferase (NnOMT6) exhibited catalytic promiscuity, mediating the methylation of diverse backbones, including phenylpropanoids, 1-benzylisoquinolines, aporphines, and protoberberines ^22^. Although several studies have been reported on the enzymes involved in the BIA biosynthetic pathway, the BIAs pathway in *N. nucifera* remains largely elusive.

Methyltransferases play a pivotal role in the biosynthesis of BIA intermediates and end products, contributing to the remarkable structural and functional diversity of BIAs^23^. Dozens of OMTs involved in BIA biosynthetic pathway have been isolated from several species in the order Ranunculales and *Liriodendron* belonging to the order Magnoliales ^23^. These OMTs primarily catalyzed the methylation of three BIA scaffolds: 1-benzylisoquinolines, protoberberine, and phthalideisoquinolines, including norcoclaurine 6-O-methyltransferase (6OMT), 3′-hydroxy-N-methylcoclaurine 4′-O-methyltransferase (4′OMT), reticuline 7-O-methyltransferase (7OMT), norreticuline 7-O-methyltransferase (N7OMT), scoulerine-9-O-methyltransferase (SOMT), scoulerine 2-O-methyltransferase (S2OMT), columbamine O-methyltransferase (CoOMT), and 4′-O-desmethyl-3-O-acetylpapaveroxine 4′-O-methyltransferase (PsOMT2:PsOMT3 and PsOMT2:Ps6OMT heterodimers) ^24,25^. However, no OMTs that catalyze the bisbenzylisoquinoline scaffold have been reported to date.

In the present study, we isolated and characterized a novel promiscuous *O*-methyltransferase, designated as Nn7OMT, which was involved in bisBIA biosynthesis in *N. nucifera*. We found that Nn7OMT catalyzed the methylation of bisbenzylisoquinoline, 1-benzylisoquinoline, aporphine, and protoberberine skeletons. We further investigated Nn7OMT function in planta by analyzing the correlation between *Nn7OMT* expression and BIA accumulation in different *N. nucifera* tissues. Additionally, we engineered the key residues of Nn7OMT via semi-rational design and obtained two mutants with enhanced substrate specificity.

## RESULTS

### Isolation and phylogeny of Nn7OMT

We performed whole-genome deep resequencing (30× coverage) for 271 representative lotus accessions collected nationwide, generating a total of 9.37 terabases (Tb) of raw data. From this dataset, we identified 22,547,166 single nucleotide polymorphisms (SNPs) and 3,582,036 insertions/deletions (InDels). Sequences from these accessions were aligned to the *N. nucifera* “China Antique” reference genome (817.9 Mb) ^26^. To investigate the genetic control of natural variation in the BIAs of lotus, the metabolite profiling data and high-quality SNPs with minor allele frequency (MAF) ≥5% were obtained and a metabolite-based genome-wide association study (mGWAS) was performed by a gene-based analysis. We identified a significant locus in controlling the accumulation of isoliensinine, which maps to chromosome 1 and exhibits a robust association with isoliensinine levels.

In searching for candidate genes underlying this locus, we found an annotated O-methyltransferase encoding gene within the confidence intervals (Fig. 1).

**Figure 1.**
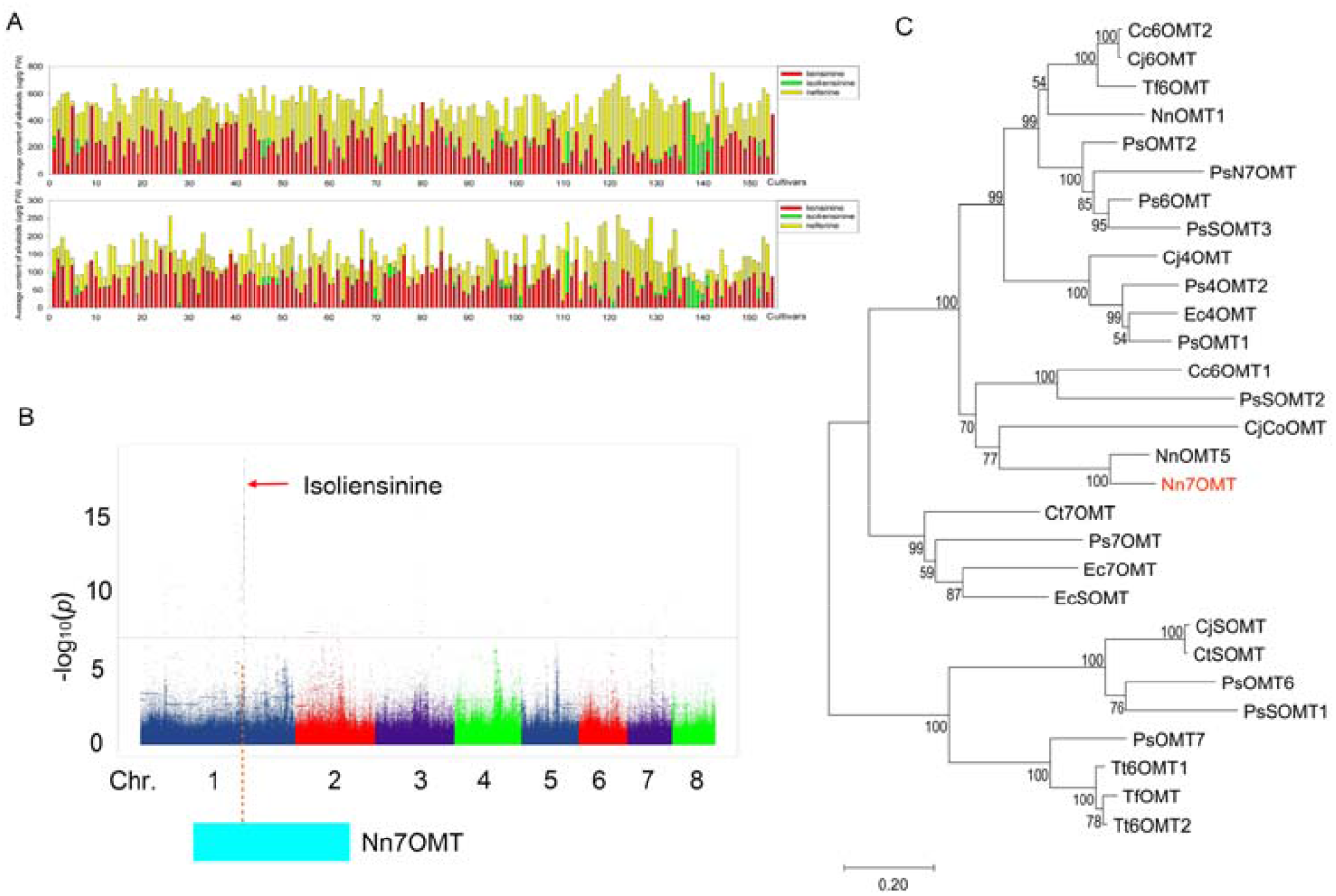
(A) The average content (ug/g fresh weight) and the proportion (%) of each alkaloid compound in 155 cultivars of lotus plumule. The numbers (x-axis) in the figure represent sample numbers. (B) Manhattan plot displaying the GWAS result of the content of isoliensinine. (C) Phylogenetic relationships between Nn7OMT and known BIA OMTs. The tree was constructed using the neighbor-joining method in MEGA7, with 1000 bootstrap replicates.

This gene was designated Nn7OMT, and its open reading frame (ORF) spans 1041 bp, encoding a 346-residue protein with a theoretical molecular weight of 37.96 kDa. Phylogenetic relationships among Nn7OMT with functionally characterized BIA OMTs was constructed to assess the potential function of Nn7OMT (Fig. 1C). Nn7OMT clustered with NnOMT5 from *N. nucifera* in a single clade, sharing 85% amino acid sequence identity. As NnOMT5 acts as a canonical 7-O-methyltransferase (7OMT) that catalyzed 7-O-methylation of the 1-benzylisoquinoline scaffold, therefore Nn7OMT was speculated to possess 7-O-methyltransferase activity.

### Molecular cloning and functional characterization of Nn7OMT

Nn7OMT candidate genes were synthesized and cloned into a pET28a vector, then expressed in Escherichia coli BL21 (DE3) strains. The enzyme was purified by Ni-NTA affinity chromatography and was analysed by SDS-PAGE. To characterize the catalytic activity of Nn7OMTs in vitro, 12 potential BIA substrates, including 6 monobenzylisoquinoline (**1**–**6**), 3 aporphine (**7**–**9**), and 3 bisbenzylisoquinoline (**10**– **12**) alkaloids, were used as substrates (Fig. S1). S-adenosyl-L-methionine (SAM) was used as the methyl donor. The reaction was analyzed using UPLC–ESI–QToF– MS/MS. Heat-inactivated enzymes (100°C, 10 min) were used as negative controls. UPLC–MS analysis showed that Nn7OMT exhibited a differential OMT catalytic activity for **1, 2, 3, 9** and **11** (Fig. 2A-E). Nn7OMT effectively catalyzed the 7-O-methylation of isoliensinine (**11**) to yield neferine (**12**), which was identified by comparing the products with a reference standard. Nn7OMT also catalyzed the C6 position of norcoclaurine (**1**) to produce coclaurine (**2**), mediated 7-O-methylation of (S)-coclaurine (**2**) and N-MethylCoclaurine (**3**) to yield (S)-norarmepavine (**4**) and armepavine (**5**), respectively. Nn7OMT catalyzed the 7-O-methylation of lirinidine (**9**) to yield nuciferine.

**Figure 2.**
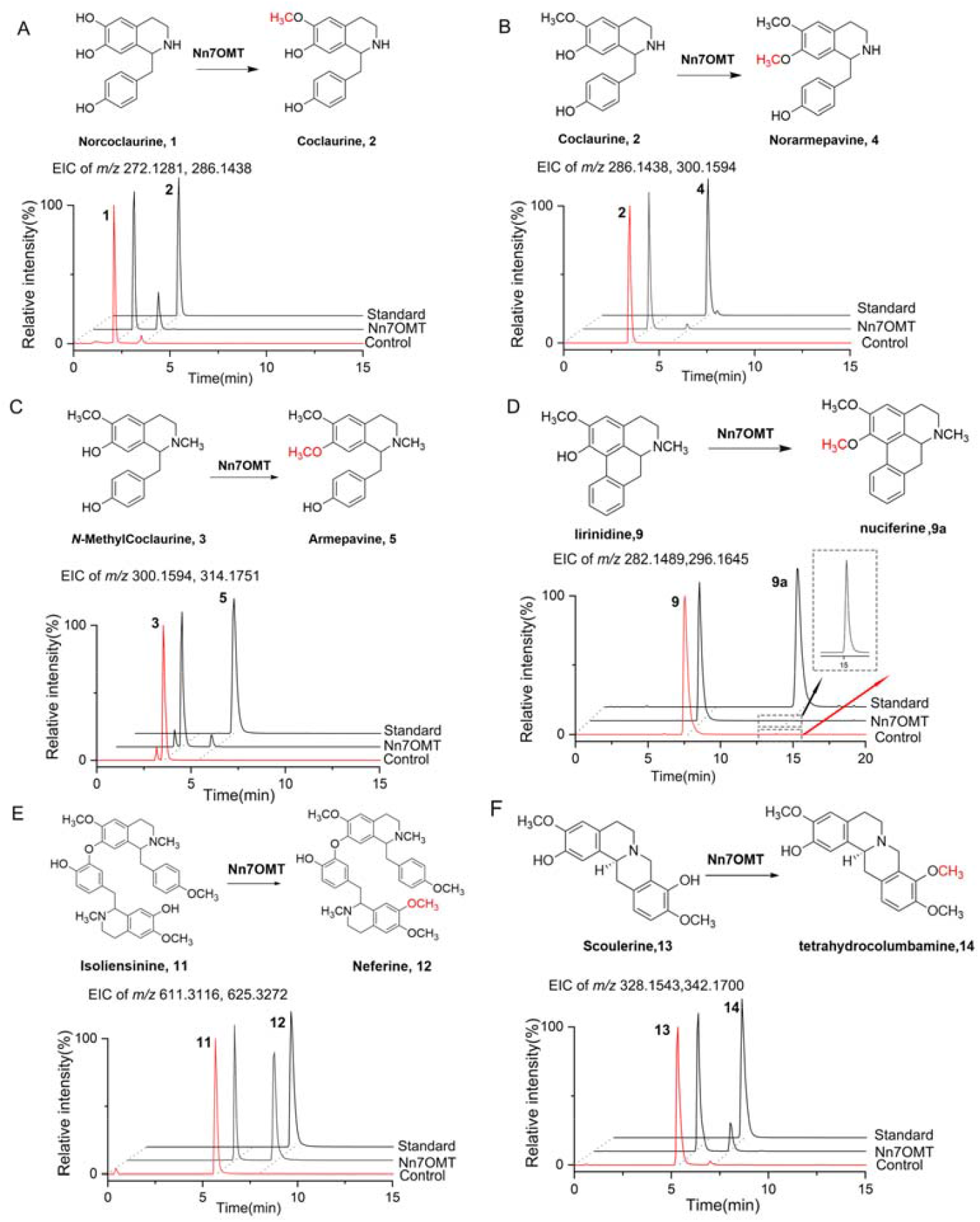
*O*-methylation of compound 1, 2, 3, 9, 11, 13 catalyzed by Nn7OMT. **(A)**The reaction and the the EIC chromatograms of *m/z* 272.1281 and 286.1438 with norcoclaurine (**1**) as the substrate. **(B)** The reaction and the EIC chromatograms of *m/z* 286.1438 and 300.1594 with coclaurine **(2)** as the substrate. **(C)** The reaction and the the EIC chromatograms of *m/z* 300.1594 and 314.1751 with *N*-methylcoclaurine **(3)** as the substrate. **(D)** The reaction and the EIC chromatograms (EIC) of *m/z* 282.1489 and 296.1645 with lirinidine **(9)** as the substrate. **(E)** The reaction and the EIC chromatograms (EIC) of *m/z* 611.3116 and 625.3272 with isoliensinine **(11)** as the substrate. **(F)** The reaction and the EIC chromatograms (EIC) of *m/z* 328.1543 and 342.1700 with scoulerine **(13)** as the substrate.

To further explore the substrate promiscuity of Nn7OMT, three protoberberine (**13, 14, 15**) substrates were used. Result showed that Nn7OMT catalyzed the methylation at the C9 position of scoulerine (**13**) to generate tetrahydrocolumbamine (Fig. 2F). All the above reaction products were identified and confirmed by comparison with reference substances. These results suggested that Nn7OMT preferentially catalyzed the methylation of the 7-hydroxyl group in BIAs. It exhibited significantly higher catalytic activity toward the bisbenzylisoquinoline alkaloid (isoliensinine) than toward the monobenzylisoquinoline alkaloids (norcoclaurine, coclaurine, and N-methylcoclaurine). Meanwhile, only negligible catalytic activity (<1%) was detected against the aporphine-type substrate (lirinidine). Notably, Nn7OMT also catalyzed the methylation at the C9 position of the protoberberine skeleton (scoulerine).

To our knowledge, Nn7OMT represents the first reported enzyme with catalytic activity toward bisbenzylisoquinoline alkaloid substrates, thereby characterizing a novel enzymatic function. As a major bioactive component in Nelumbo nucifera, neferine exerts diverse pharmacological effects, including hypoglycemic, hypotensive, sedative, and antiarrhythmic activities. Given its catalytic profile, Nn7OMT holds great potential as a biocatalyst for the biosynthesis of neferine.

### Biochemical characterization of recombinant Nn7OMT protein

The enzyme characteristics of Nn7OMT were studied (Fig. 3). The pH and temperature conditions influencing the enzyme activity were optimized. Nn7OMT exhibited its maximum activity at pH 8.0 (50 mM potassium phosphate) and 37°C. The kinetic property of Nn7OMT was determined using isoliensinine (**11**) as substrate. The Km value of Nn7OMT for isoliensinine was 86.47±0.77 μM, Vmax value was 26.76±0.02 nmol·min-1·mg-1, and kcat/Km value was 195.79±1.62 M-1·s-1.

**Figure 3.**
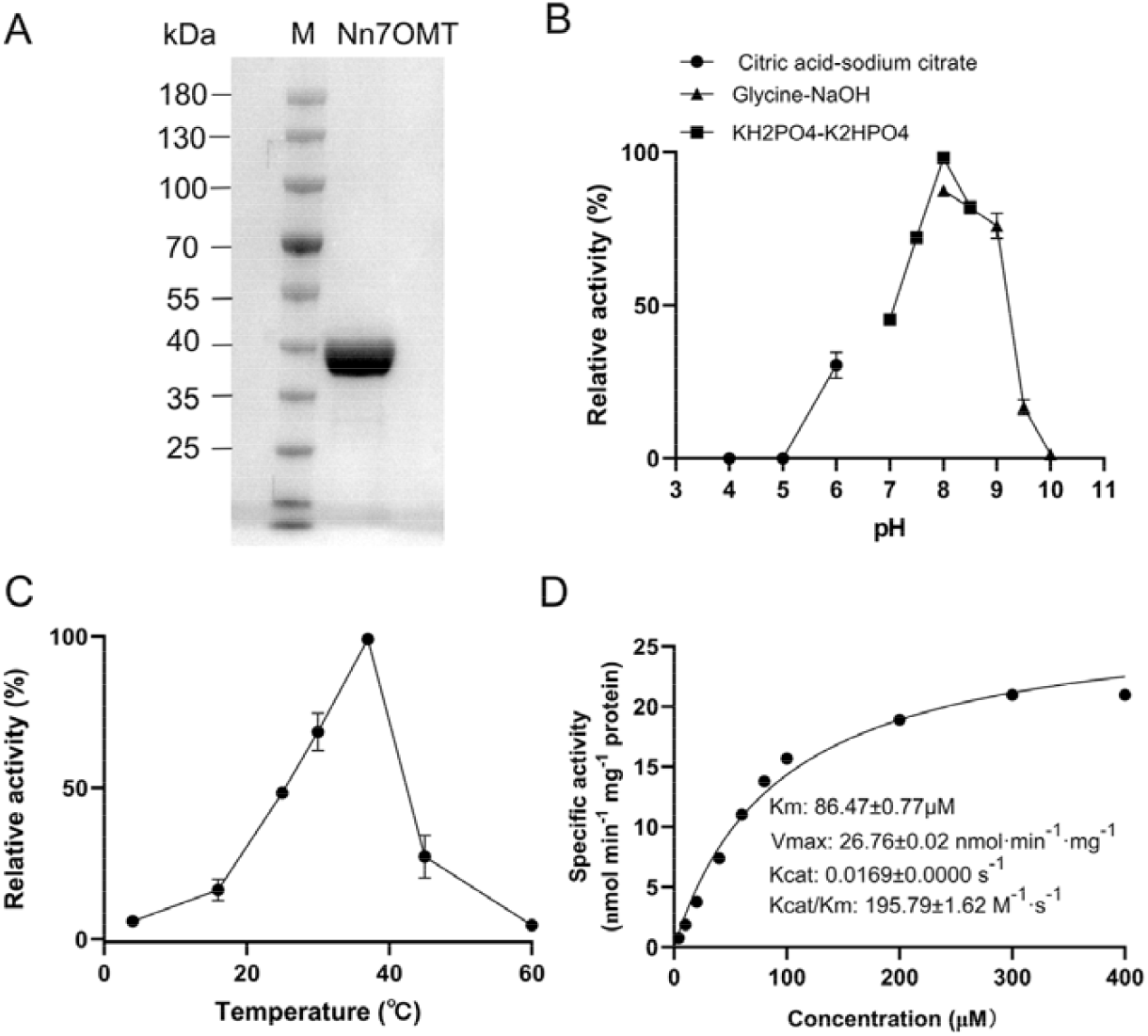
Biochemical characterization of recombinant Nn7OMT protein. (A) SDS-PAGE analysis of purified recombinant Nn7OMT. Effects of pH (B), and temperature (C) on enzyme activity of Nn7OMT. (D) The Michaelis-Menten profile of Nn7OMT using various concentrations of isoliensinine. Error bars represented standard deviation of three independent replicates.

### Nn7OMT contributes to the biosynthesis of neferine in vivo

To further clarify the functional relevance of Nn7OMT in BIA biosynthesis, we analyzed its expression pattern across four tissues (tender leaves, mature leaves, flowers, embryos) from three *N. nucifera* varieties and correlated it with bisbenzylisoquinoline alkaloid accumulation (Fig. 4). Notably, Nn7OMT exhibited an extremely strong tissue-specific expression profile, being exclusively expressed in the embryos, while its expression was barely detectable in other tested tissues (e.g., tender leaves, mature leaves, and flowers). Consistently, three bisbenzylisoquinoline alkaloids were predominantly accumulated in the embryo, which showed a striking consistency with the expression pattern of Nn7OMT. These findings underscore the tight correlation between Nn7OMT expression and the accumulation of bisbenzylisoquinoline alkaloids in the embryo, strongly suggesting that Nn7OMT is closely involved in the biosynthesis of neferine in planta.

**Figure 4.**
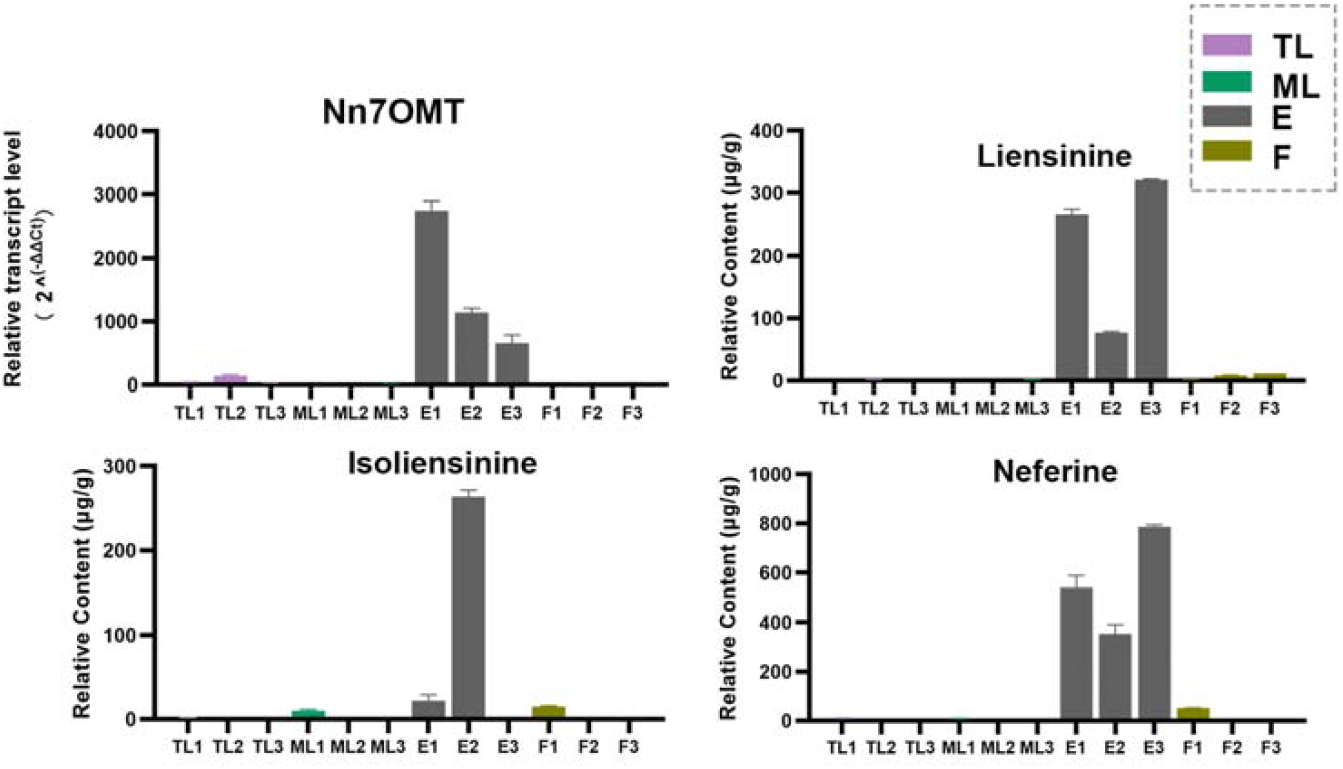
Relative expression of Nn7OMT and three bisBIA contents of different tissues in three *N*.*nucifera* cultivars. Error bars indicated standard deviation of three independent replicates. The tissue samples were listed as follows: TL1-3, tender leaves of three cultivars; ML1-3, mature leaves of three cultivars; F1-3, flowers of three cultivars; E1-3, embryos of three cultivars.

### Engineering key residues and site-directed mutagenesis of Nn7OMT

To further explore the catalytic mechanisms of Nn7OMT enzyme, the protein structure of Nn7OMT was de novo modeling using Alphafold2 (Fig. S2). Then, the substrate isoliensinine was docked into the structure of Nn7OMT using Molecular Operating Environment (MOE) dock. To search for the stable structure of Nn7OMT-SAH-Isoliensinine, we performed a molecular docking search followed by all-atom, explicit water MD simulations (Fig.5 A-B). The result that root means square deviation (RMSD) of the backbone of Nn7OMT is less than 4.13 Å and that of Isoliensinine is less than 2.67 Å (Fig. S3A). The system achieves equilibrium within the simulation time, which suggest that the force field and simulation protocol are adequate. Root mean square fluctuation (RMSF) can indicate the flexibility of amino acid residues in the protein (Fig. S3B). The residue Glu141, Phe144 and Ser303 in Nn7OMT formed hydrogen bond with Isoliensinine, and the residue Asp232, Leu213, Asp212, Ser191, Lys246, Asp187, Val188 and Met233 formed hydrogen bond with SAH (Fig.5 C-D). To further identify the key active residues of Nn7OMT, the binding energy (ΔGtotal) of Isoliensinine with complex Nn7OMT-SAH was calculated using the MM-PBSA method. The binding free energy (ΔGtotal) was composed of Van der Waals (ΔEvdw), electrostatic interactions (ΔEele), the polar solvation (ΔGpolar) and nonpolar solvation (ΔGnonpolar) Energy. The binding free energy upon isoliensinine with complex Nn7OMT-SAH was computed to be −54.58 kcal/mol in aqueous environments (Tab.S1). The Nn7OMT-SAH-Isoliensinine binding was largely governed by Van der Waals (ΔEvdw). Further energy composition analysis suggests that residues Pro308, Leu154, Thr307, Leu140, MET158, Leu109, Phe144, Val306, Ser191 and Met304 in Isoliensinine contribute most to the complex binding free energy (Tab.S2).

In order to probe the catalytic mechanism, the key amino acid residues were mutated to alanine, and the catalytic activity of mutants was screened using crude proteins with six compounds (**1, 2, 3, 11**) as substrates. The catalytic activity of all mutants showed no significant difference to with wild Nn7OMT for isoliensinine substrate (Fig. 5E). However, most of mutants showed remarkably decreased catalytic activities for 1-benzylisoquinoline alkaloids (**1, 2, 3**), especially mutants P308A, T307A, M158A and V306A. The mutant P308A significantly decreased the activity for coclaurine (**2**), and mutant L154A significantly decreased the activity of N-methylcoclaurine (**3**), which indicated that residues P308 and L154 plays a key role in the activity of Nn7OMT to substrates **2** and **3**, respectively. The mutants M158A and V306A have hardly activity for **1, 2, 3**. Although, there was no mutant to remarkablely improve catalytic activity for Nn7OMT, mutants M158A and V306A maintained catalytic activity for **11** and simultaneously resulted in hardly catalytic activity with 1-benzylisoquinoline alkaloids (**1, 2, 3**) substrates. The result improved the substrate specificity of Nn7OMT, which contributes to reducing by-product formation and purification difficulty in its practical application as a biosynthesis biocatalyst.

**Figure 5.**
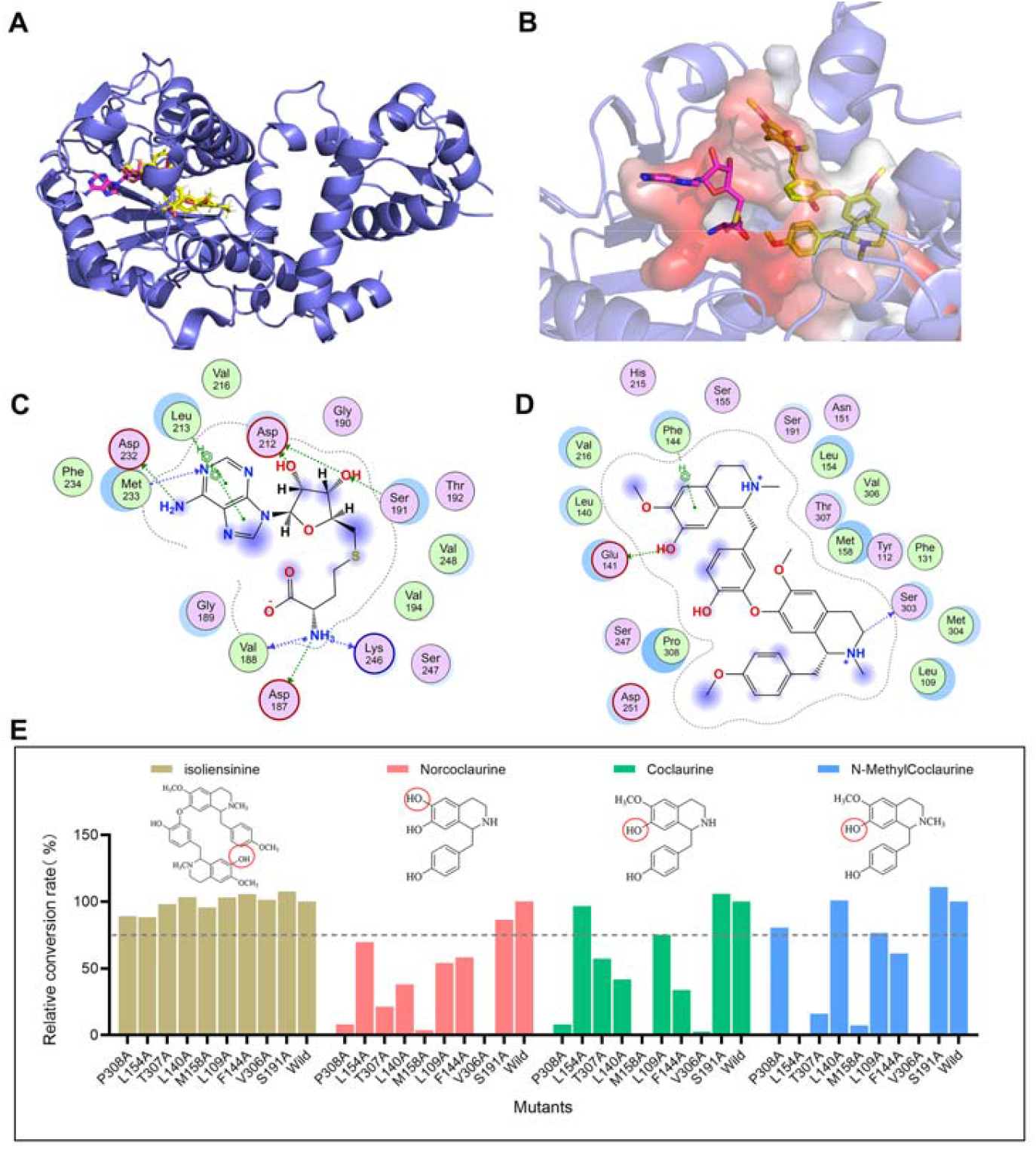
Semi-rational design and mutants screening for Nn7OMT. (A) The binding model of complex Nn7OMT-SAH-Isoliensinine. (B) Substrate binding pockets of complex Nn7OMT-SAH-Isoliensinine. The isoliensinine was colored in yellow, the SAH was colored in magenta, and the Nn7OMT was colored in lightblue. key residues surrounding SAH (C) and isoliensinine (D) in the active site of Nn7OMT. (E) Functional screening of the mutants with compounds **1, 2, 3**, and **11** as the substrate.

## DISCUSSION

Benzylisoquinoline alkaloids (BIAs) from *N. nucifera* displayed distinct structural and stereochemical characteristics relative to those of Ranunculales species^27,28^. Several bisbenzylisoquinoline alkaloids (bisBIAs), such as liensinine, isoliensinine, and neferinine, were lotus-specific metabolites. These compounds exerted notable pharmacological activities, including anticancer, hypoglycemic, hypotensive, sedative, and antiarrhythmic effects ^5,29,30^. However, the molecular mechanisms governing bisBIA biosynthesis in *N. nucifera* remain unclear. Methylation played a pivotal role in the functionalization of specialized metabolites. Although ∼50 O-methyltransferases (OMTs) involved in BIA biosynthesis have been functionally characterized from plants of Ranunculales, Magnoliales, and Proteales, none has been reported to catalyze bisBIA formation ^23^. In this study, we characterized a novel O-methyltransferase (OMT) from *Nelumbo nucifera*, designated Nn7OMT, which mainly catalyzed the bisBIA skeleton isoliensinine to produce neferinine. Notably, Nn7OMT represents the first OMT shown to catalyze the bisBIA backbone, thus highlighting its critical significance for neferinine biosynthesis.

Fifteen BIA substrates with diverse structural backbones were employed to screen the catalytic activity of Nn7OMT, including six 1-benzylisoquinolines, three aporphines, three bisBIAs, and three protoberberines. Nn7OMT exhibited catalytic promiscuity, catalyzing O-methylation at the C6/C7 positions of the 1-benzylisoquinolines norcoclaurine, coclaurine and N-methylcoclaurine, the C7 position of the aporphine lirinidine, and the C9 position of the protoberberine scoulerine, with isoliensinine identified as its preferred substrate. The expression profile of Nn7OMT was in consistent agreement with bisBIA accumulation in lotus plants, where bisBIAs are exclusively accumulated in embryonic tissues and only trace amounts detectable in all other vegetative tissues. This finding thus further substantiates the physiological function of Nn7OMT in the planta biosynthesis of bisBIAs. The novel catalytic activity of Nn7OMT toward bisBIAs highlights the evolutionary divergence of the BIA metabolic pathway in *N. nucifera*, and further underscores the species-specific capacity of this plant to synthesize specialized bisBIA alkaloids. Our findings thus greatly advance the current understanding of the biosynthetic pathway underlying bisBIA production in *N. nucifera*.

To further elucidate the catalytic mechanism of Nn7OMT, key residues that predominantly contribute to substrate binding and catalysis were identified by AlphaFold2 de novo modeling, MOE molecular docking, MD simulations, and MM-PBSA energy decomposition. The functional roles of these residues were validated using site-directed mutagenesis. Alanine mutation of 9 critical residues (Pro308, Leu154, Thr307, Leu140, Met158, Leu109, Phe144, Val306, Ser191) identified by MM-PBSA showed no significant change in catalytic activity toward isoliensinine, but resulted in a marked decrease in activity toward 1-benzylisoquinoline alkaloids (**1, 2, 3**). Specifically, P308A impaired activity toward coclaurine (**2**), L154A diminished activity toward N-methylcoclaurine (**3**), while Met158A and Val306A almost completely lost catalytic activity toward substrates **1, 2**, and **3**. These results indicated that these residues play crucial roles in the catalytic activity of Nn7OMT toward 1-benzylisoquinoline alkaloids, possibly by regulating substrate binding stability or the stereo-spatial distance between the methyl donor SAH and substrate acceptors. Interestingly, Met158A and Val306A retained catalytic activity toward compound **11** while losing activity toward 1-benzylisoquinoline alkaloids, representing a notable improvement in substrate specificity. Furthermore, this engineered specificity effectively reduces by-product formation from non-target substrates and lowers purification difficulty. Collectively, these results demonstrate that Met158A and Val306A mutants hold potential as biocatalysts for the synthesis of specialized bisBIAs in synthetic biology platforms, given their tailored substrate selectivity.

## MATERIALS AND METHODS

### Plant materials and chemicals

Standards norcoclaurine (**1**), coclaurine (**2**), N-methylcoclaurine (**3**), norarmepavine (**4**), armepavine (**5**), lotusine (**6**), neferine (**12**) were supplied from Shanghai Yuanye Biotech (Shanghai, China). asimilobine (**7**), and *N*-methlyasimilobine (**8**), lirinidine (**9**), liensinine (**10**), and isoliensinine (**11**), scoulerine **(13)**, tetrahydrocolumbamine **(14)**, jatrorrhizine **(15)** were purchased from Chengdu Push Biotech (Chengdu, China); SAM was derived from Sigma-Aldrich (St. Louis, MO, USA). BeyoGoldTM His-tag Purification Resin was purchased from Beyotime Biotech (Shanghai, China). Acetonitrile, methanol, and formic acid used in UPLC-MS/MS analysis were obtained from Sigma-Aldrich. Different tissues of the *N. nucifera* cultivar “Hongtailian” were collected and immediately frozen in liquid nitrogen and stored at −80°C until use.

### Isolation and phylogenetic analysis of Nn7OMT candidate genes

Raw resequencing data of *N. nucifera* were generated in our preliminary work. All these data were aligned to the reference genome of *N. nucifera* “Ancient Chinese Lotus” following the standard bwa-GATK pipeline, to call single nucleotide polymorphism (SNP) loci and their corresponding genotypic information. GWAS was performed with FaST-LMM software: representative SNPs with MAF > 5% and missing sample rate < 10% were selected, and a mixed linear model (MLM) incorporating population structure was applied to calculate the correlation *P*-values between all SNPs and each metabolite. The significance threshold of *P*-values was determined as 0.05/*n* and further corrected by Bonferroni correction. For significantly associated SNPs, the maximum variation explained by linked SNPs and the explained variation per individual SNP were calculated for each metabolite. The distribution of significant SNPs was analyzed in 1-Mb genomic windows, and 1000 iterations were performed for SNPs with *P* < 0.01 to estimate genetic hotspots. Candidate genes were finally inferred based on coding region variations associated with extreme metabolite contents and their functional annotation results.

We utilized ClustalW to align the amino acid sequences and employed MEGA 7.0 software to construct a phylogenetic tree for Nn7OMT and the 28 other OMTs with 1000 bootstrap replicates.

### Molecular cloning, expression and enzyme activity assays of recombinant Nn7OMTs

The coding region (CDS) of Nn7OMT was synthesized by Tsingke Biotechnology Co., Ltd. and inserted into a pET-28a expression vector using a ClonExpress® II One Step Cloning Kit (Vazyme Biotech, Nanjing, China). The recombinant plasmids were transformed into E. coli BL21 (DE3) (TransGen Biotech, China) for expressing protein. The cells were cultured in 300 mL of LB medium (50 μg/mL kanamycin) at 37°C and shaken at 200 rpm until OD600 reached 0.6–0.8. After that, the cultures were added a final concentration of 0.3 mM IPTG and induced at 16°C for 20 h with shaking (180 rpm). The cell pellets were harvested at 4°C by centrifugation and then resuspended with lysis buffer (50 mM NaH2PO4, 300 mM NaCl, 20 mM imidazole, 2% glycerol, pH 8.0). Thereafter, the cells of suspension were disrupted via sonication for 30 min in an ice bath, and centrifuged at 10000 rpm for 40 min at 4°C. The supernatant was subsequently added to 2 mL of pre-equilibrated BeyoGold™ His-tag Purification Resin (Beyotime Biotech, China) and incubated at 4°C for 4 h. Thereafter, the lysis buffer and the elution buffers containing 250 mM imidazole were used to wash and elute the protein, respectively. The purity of recombinant protein was examined by SDS–PAGE analysis (10% gel) and concentrated using a 10KDa ultrafiltration tube (Merck Millipore). The concentration of purified enzyme was determined with Bradford Protein Assay kit (TransGen Biotech, Beijing, China).

The catalytic activity of recombinant Nn7OMT proteins was investigated using 12 potential BIA substrates containing monobenzylisoquinolines 1–6, aporphine alkaloids 7–9, and bisbenzylisoquinolines 10–12. The reaction was conducted in a 100-µL mixture system comprising 50 mM potassium phosphate (pH 8.0), 200 µM SAM donor, 200 µM acceptor substrate, and 50 μg purified protein. Heat-inactivated enzymes (100 °C, 10 min) were used as negative controls. The mixtures were incubated at 37°C for 12 h, terminated by the addition of 100 µL methanol, then centrifuged at 12,000 ×g for 15 min at 4°C before UPLC–MS/MS analysis.

### Biochemical characterization of Nn7OMT

The reaction time, pH and temperature of Nn7OMT were optimized using isoliensinine **(11)** as substrate. Enzye assays were performed in 50 mM Citric acid-sodium citrate buffer (pH 4.0-6.0), 50 mM potassium phosphate buffer (pH 7.0-8.5) and 100 mM Gly–NaOH buffer (pH 8.0-10.0) respectively at 37°C for 4 h to obtain the optimal pH. The reactions were conducted at 4°C–60°C for 4 h (pH 8.0) to optimize temperature. The kinetic parameters of Nn7OMT were determined in a 100-µL reaction mixture containing 50 mM potassium phosphate (pH 8.0) and varying concentrations of isoliensinine (4–400 µM) at a fixed concentration of 200 µM SAM. The reactions were incubated at 37°C for 20 min and terminated by adding 100 µL methanol. All reactions were conducted in triplicates. The kinetics datas were calculated by nonlinear regression analysis using a Michaelis–Menten model in the GraphPad Prism 8.0 software.

### Gene expression and alkaloid content analysis in different organs

The extraction of total RNA and synthesis of cDNAs from four tissues (tender leaves, mature leaves, flowers, and embryos) in three *N. nucifera* varieties were performed as previously described. Expression levels of Nn7OMT were determined by reverse transcription quantitative PCR (RT-qPCR) analysis with Taq Pro Universal SYBR qPCR Master Mix (Vazyme Biotech). Primer sequences for RT-qPCR were presented in Tab. S3. β-actin was used as the endogenous reference gene to calculate the relative transcript abundance. All experiments were performed with three replicates.

The method for extracting alkaloids from different samples was same to that described in our previous study ^22^. Fine powder samples (100 mg) were weighed carefully, extracted using 1 mL of a mixture of 70 % methanol and 30 % water (v:v), agitated six times for ∼30 s in intervals of 30 min, and then incubated at 4 °C overnight. After centrifuging the samples for 10 min at 12,000 rpm, the supernatants were filtered through 0.22-µm filters before further UPLC–ESI–QTOF–MS/MS analysis.

### De novo modeling, molecular docking, and MD simulation

De novo modeling of protein Nn7OMT was conducted in Alphafold2. MOE2 Dock was used for molecular docking of Nn7OMT-SAH with Isoliensinine. The Nn7OMT-SAH complex was obtained by superposition of the SAH on the crystal structure of Tf6OMT (PDB ID: 5ICE) into the protein Nn7OMT, and binding site of native ligand in template structure was set as binding pocket for isoliensinine.

After docking, the complex Nn7OMT-SAH–isoliensinine was optimized using MD simulation performed with AMBER16. To neutralize protein structure, sodium/chlorine counterions were added. The solvation of each protein structure was conducted in a cuboid box of TIP3P water molecules with a 10 Å solvent layers between the edges of the box and the surface of the solute. The protein was subjected to AMBER GAFF and FF14SB force fields. All covalent bonds involving hydrogen atoms were restricted using the SHAKE algorithm with a time step of 2 FS. The long-range electrostatic interactions were treated by Particle mesh Ewald method. Each solvated system was performed two steps of minimization before the heating step. The initial minimization was conducted with 4000 cycles with all heavy atoms restrained using 50 kcal/(mol·Å2), whereas solvent molecules and hydrogen atoms were free to move. Then, the stage of non-restrained minimization was performed with the steepest descent minimization for 2000 cycles and conjugated gradient minimization for 2000 cycles. Thereafter, the entire system was heated from 0 K to 300 K in 100 ps using Langevin dynamics at a constant volume, following which it was equilibrated for 150 ps at a constant pressure of 1 atm. Periodic boundary dynamics simulations were performed for the entire system in an NpT ensemble (constant composition, pressure, and temperature) at a constant pressure of 1 atm and 300 K during the production step. In production phase, 100-ns simulation was conducted. The binding free energy of the complex was calculated using the MM/PBSA method.

### Site-directed mutagenesis of Nn7OMT

The mutant plasmids of Nn7OMT were cloned with pET28a-Nn7OMT vector as the template using a Fast MultiSite Mutagenesis System kit (TransGen, China). Primer pairs used to PCR amplification were showed in Tab. S4.

### UPLC–ESI–QToF–MS/MS conditions

Determining the content of BIAs and enzyme assay products was performed as described previously ^31^. An Agilent 1290 photodiode array and a 6540 triple quad mass time-of-flight mass spectrometer were used for the UPLC-QTOF-MS/MS analysis using a dual electrospray ionization (ESI) detector. A Waters ACQUITY UPLC CSH C18 Column (2.1 mm × 100 mm, 1.7 μm) was used to separate and analyze the samples with a mobile phase of 0.1% formic acid (eluent A) and acetonitrile (eluent B). The following elution gradient was used: 2% B at 0 min, 5% B at 1 min, 9% B at 5 min, 10% B at 12 min, 15% B at 16 min, 45% B at 20 min, 100% B at 22 min; flow rate, 0.3 mL/min; injection volume, 2 μL; and temperature, 35°C.

The ESI source was set to positive ionization mode and the following parameters were set for the QTOF-MS detector: nebulizer, 45 psig; nozzle voltage, 500 V; Vcap, 4000 V; cone voltage, 20 V; sheath gas temp, 350°C; drying gas, 8 L min^−1^; sheath gas flow, 11 L min^−1^; collision energy, 30 eV and scan range, *m/z* 100–1500 Da. The measured masses were modified by internal references (purine and HP-0921) in real-time.

## CRediT authorship contribution statement

Yuetong Yu, Sha Chen - designed the research; **Yuetong Yu, Zhennan Wang**, and **Sha Chen -** performed the research; **Yuetong Yu, Xinyi Qi-** collected the samples; **Yuetong Yu** and **Sha Chen** - analyzed the data and drafted the paper; **Yuetong Yu** and **Sha Chen** revised the paper. **Yuetong Yu, Zhennan Wang**, and **Sha Chen -** revised the paper.

## Data availability statement

All data supporting this research can be obtained in the paper and within its supplementary materials published online.

## Acknowledgements

This work was supported by the National Natural Science Foundation of China (32170388), Scientific and Technological Innovation project of China Academy of Chinese Medical Sciences (CACMS Innovation Fund CI2024E003XY-18), Fundamental Research Funds for the Central public welfare research institutes of China (ZXKT25026, ZXKT25043, ZZ13-YQ-057), and Fundamental Research Program of Shanxi Province (202403021212244).

## Supporting Information statement

Supplementary data to this article can be found online.

## References

1 Chen, G., Zhu, M. & Guo, M. Research advances in traditional and modern use of Nelumbo nucifera: phytochemicals, health promoting activities and beyond. Crit Rev Food Sci Nutr 59, S189–s209, doi:10.1080/10408398.2018.1553846 (2019).

2 Yang, H. et al. Lotus (Nelumbo nucifera): a multidisciplinary review of its cultural, ecological, and nutraceutical significance. Bioresources and bioprocessing 11, 18, doi:10.1186/s40643-024-00734-y (2024).

3 Zheng, H. et al. Research Advances in Lotus Leaf as Chinese Dietary Herbal Medicine. Am J Chin Med 50, 1423–1445, doi:10.1142/s0192415x22500616 (2022).

4 Zhao, X. et al. Recent advances on bioactive compounds, biosynthesis mechanism, and physiological functions of Nelumbo nucifera. Food Chem 412, 135581, doi:10.1016/j.foodchem.2023.135581 (2023).

5 Wang, Z. et al. Alkaloids from lotus (Nelumbo nucifera): recent advances in biosynthesis, pharmacokinetics, bioactivity, safety, and industrial applications. Crit Rev Food Sci Nutr 63, 4867–4900, doi:10.1080/10408398.2021.2009436 (2023).

6 Kashiwada, Y. et al. Anti-HIV benzylisoquinoline alkaloids and flavonoids from the leaves of Nelumbo nucifera, and structure-activity correlations with related alkaloids. Bioorg Med Chem 13, 443–448, doi:10.1016/j.bmc.2004.10.020 (2005).

7 Liu, C. P. et al. Inhibition of (S)-armepavine from Nelumbo nucifera on autoimmune disease of MRL/MpJ-lpr/lpr mice. Eur J Pharmacol 531, 270–279, doi:10.1016/j.ejphar.2005.11.062 (2006).

8 Ka, S. M. et al. (S)-armepavine from Chinese medicine improves experimental autoimmune crescentic glomerulonephritis. Rheumatology (Oxford) 49, 1840–1851, doi:10.1093/rheumatology/keq164 (2010).

9 Qi, Q. et al. Identification of the anti-tumor activity and mechanisms of nuciferine through a network pharmacology approach. Acta Pharmacol Sin 37, 963–972, doi:10.1038/aps.2016.53 (2016).

10 Wan, Y. et al. Nuciferine, an active ingredient derived from lotus leaf, lights up the way for the potential treatment of obesity and obesity-related diseases. Pharmacol Res 175, 106002, doi:10.1016/j.phrs.2021.106002 (2022).

11 He, C. L. et al. Identification of bis-benzylisoquinoline alkaloids as SARS-CoV-2 entry inhibitors from a library of natural products. Signal transduction and targeted therapy 6, 131, doi:10.1038/s41392-021-00531-5 (2021).

12 Xu, L. et al. Neferine induces autophagy of human ovarian cancer cells via p38 MAPK/JNK activation. Tumour Biol 37, 8721–8729, doi:10.1007/s13277-015-4737-8 (2016).

13 Wu, C. et al. Mitochondrial protective effect of neferine through the modulation of nuclear factor erythroid 2-related factor 2 signalling in ischaemic stroke. Br J Pharmacol 176, 400–415, doi:10.1111/bph.14537 (2019).

14 Desgagne-Penix, I. & Facchini, P. J. Systematic silencing of benzylisoquinoline alkaloid biosynthetic genes reveals the major route to papaverine in opium poppy. Plant J 72, 331–344, doi:10.1111/j.1365-313X.2012.05084.x (2012).

15 W, B. G. A. & J, F. P. Benzylisoquinoline alkaloid biosynthesis in opium poppy. Planta 240 (2014).

16 Minami, H. Fermentative production of plant benzylisoquinoline alkaloids in microbes. Bioscience, biotechnology, and biochemistry 77, 1617–1622, doi:10.1271/bbb.130106 (2013).

17 Singh, A., Menéndez-Perdomo, I. M. & Facchini, P. J. Benzylisoquinoline alkaloid biosynthesis in opium poppy: an update. Phytochemistry Reviews 18, 1457–1482, doi:10.1007/s11101-019-09644-w (2019).

18 Vimolmangkang, S. et al. Evolutionary origin of the NCSI gene subfamily encoding norcoclaurine synthase is associated with the biosynthesis of benzylisoquinoline alkaloids in plants. Sci Rep 6, 26323, doi:10.1038/srep26323 (2016).

19 Hao, C., Yu, Y., Liu, Y., Liu, A. & Chen, S. The CYP80A and CYP80G Are Involved in the Biosynthesis of Benzylisoquinoline Alkaloids in the Sacred Lotus (Nelumbo nucifera). International Journal of Molecular Sciences 25, doi:10.3390/ijms25020702 (2024).

20 Pyne, M. E., Gold, N. D. & Martin, V. J. J. Pathway elucidation and microbial synthesis of proaporphine and bis-benzylisoquinoline alkaloids from sacred lotus (Nelumbo nucifera). Metab Eng 77, 162–173, doi:10.1016/j.ymben.2023.03.010 (2023).

21 Menendez-Perdomo, I. M. & Facchini, P. J. Isolation and characterization of two O-methyltransferases involved in benzylisoquinoline alkaloid biosynthesis in sacred lotus (Nelumbo nucifera). J Biol Chem 295, 1598–1612, doi:10.1074/jbc.RA119.011547 (2020).

22 Yu, Y. et al. Functional characterization and key residues engineering of a regiopromiscuity O-methyltransferase involved in benzylisoquinoline alkaloid biosynthesis in Nelumbo nucifera. Hortic Res 10, uhac276, doi:10.1093/hr/uhac276 (2023).

23 Morris, J. S. & Facchini, P. J. Molecular Origins of Functional Diversity in Benzylisoquinoline Alkaloid Methyltransferases. Front Plant Sci 10, 1058, doi:10.3389/fpls.2019.01058 (2019).

24 Purwanto, R., Hori, K., Yamada, Y. & Sato, F. Unraveling Additional O-Methylation Steps in Benzylisoquinoline Alkaloid Biosynthesis in California Poppy (Eschscholzia californica). Plant Cell Physiol 58, 1528–1540, doi:10.1093/pcp/pcx093 (2017).

25 Dastmalchi, M., Park, M. R., Morris, J. S. & Facchini, P. Family portraits: the enzymes behind benzylisoquinoline alkaloid diversity. Phytochemistry Reviews 17, 249–277, doi:10.1007/s11101-017-9519-z (2017).

26 Qi, H., Yu, F., Deng, J., Zhang, L. & Yang, P. The high-quality genome of lotus reveals tandem duplicate genes involved in stress response and secondary metabolites biosynthesis. Hortic Res 10, uhad040, doi:10.1093/hr/uhad040 (2023).

27 Menéndez-Perdomo, I. M. & Facchini, P. J. Benzylisoquinoline Alkaloids Biosynthesis in Sacred Lotus. Molecules 23, doi:10.3390/molecules23112899 (2018).

28 Hao, C. et al. Visualization and identification of benzylisoquinoline alkaloids in various nelumbo nucifera tissues. Heliyon 9, doi:10.1016/j.heliyon.2023.e16138 (2023).

29 Itoh, A. et al. Bisbenzylisoquinoline Alkaloids from Nelumbo nucifera. Chemical & pharmaceutical bulletin 59, 947–951, doi:10.1248/cpb.59.947 (2011).

30 Chen, S. et al. Natural alkaloids from lotus plumule ameliorate lipopolysaccharide-induced depression-like behavior: integrating network pharmacology and molecular mechanism evaluation. Food Funct 10, 6062–6073, doi:10.1039/c9fo01092k (2019).

31 Yu, Y. et al. Identification and quantification of oligomeric proanthocyanidins, alkaloids, and flavonoids in lotus seeds: A potentially rich source of bioactive compounds. Food Chem 379, 132124, doi:10.1016/j.foodchem.2022.132124 (2022).

