## Supplementary information for "Functional characterization and molecular engineering of a *O*-methyltransferase involved in bis-benzylisoquinoline alkaloids biosynthesis from *Nelumbo nucifera*"

**Figure S1.** Structures of BIAs substrate including monobenzylisoquinoline **1**– **7**, aporphine **8**–**10**, and bisbenzylisoquinoline **11**–**13**.

**Figure S2.** (A) The predicted model of Nn7OMT. (B) Ramachandran plot for Nn7OMT. Dark blue dots represent the residues in favored regions; orange dots represent the residues in allowed regions.

**Figure S3.** (A) Root means square deviation (RMSD) of Nn7OMT backbone and substrate norcoclaurine. (B) RMSF of the protein structure model in molecular dynamics simulation within 100 ns.

**Table S1.** Average Binding Energy of Nn7OMT-SAH-Isoliensinine

**Table S2.** Energy decomposition of key residues contributing most to the complex binding free energy

**Table S3.** Primers for qRT-PCR.

**Table S4.** Primers for site-directed mutagenesis of Nn7OMT.

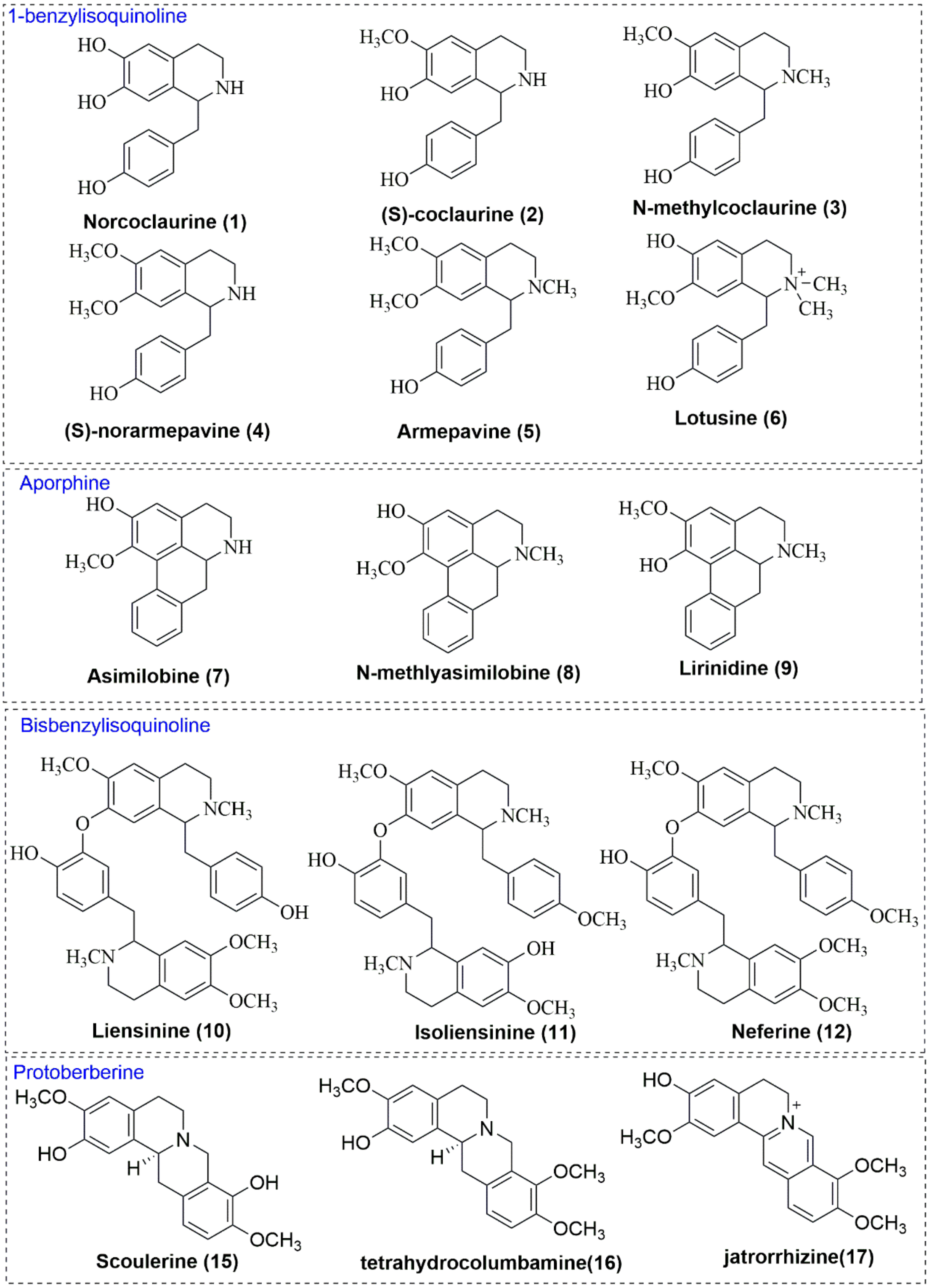
1-benzylisoquinoline
Norcoclaurine (1)
(S)-coclaurine (2)
N-methylcoclaurine (3)
(S)-norarmepavine (4)
Armepavine (5)
Lotusine (6)
Aporphine
Asimilobine (7)
N-methlyasimilobine (8)
Lirinidine (9)
Bisbenzylisoquinoline
Liensinine (10)
Isoliensinine (11)
Neferine (12)
Protoberberine
Scoulerine (15)
tetrahydrocolumbamine(16)
jatrorrhizine(17)

**Figure S1**

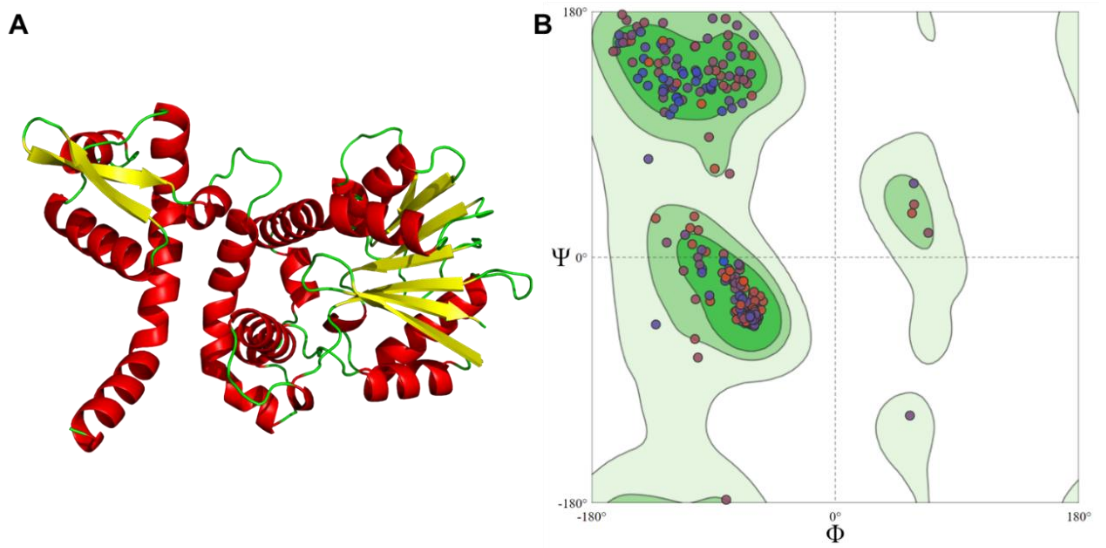


**Figure S2**

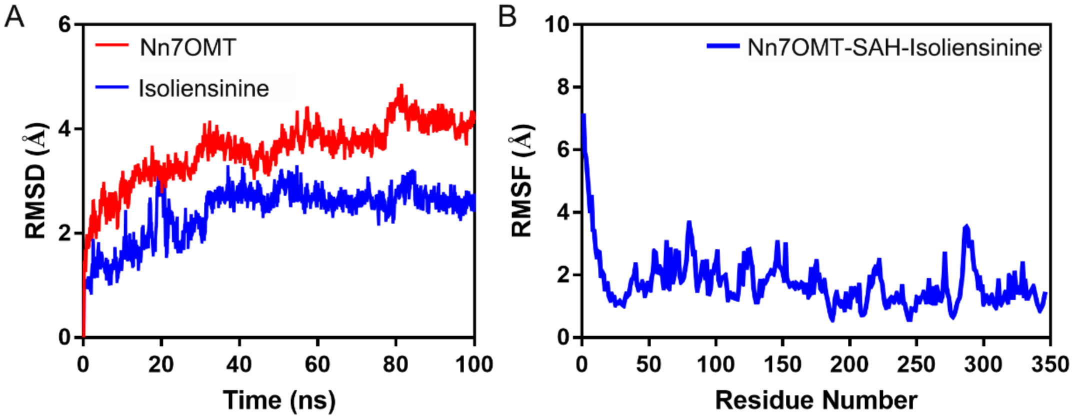


**Figure S3**

**Table S1.** Average Binding Energy of Nn7OMT-SAH-Isoliensinine

| Contribution | Energy (kcal/mol) |
|---|---|
| $\Delta E_{vdw}$ | -69.73±0.15 |
| $\Delta E_{ele}$ | -13.68±0.26 |
| $\Delta G_{polar}$ | 37.69±0.21 |
| $\Delta G_{nonpolar}$ | -8.87±0.02 |
| $\Delta G_{total}$ | -54.58±0.19 |

**Table S2.** Energy decomposition of key residues contributing most to the complex binding free energy

| Residue | ΔEvdw | ΔEele | ΔGsol | ΔGtotal |
|---|---|---|---|---|
| Pro308 | -2.74194 | -0.25784 | 0.061745 | -2.93803 |
| Leu154 | -2.37374 | 0.088703 | -0.03904 | -2.32408 |
| Thr307 | -1.64673 | -1.71184 | 1.074442 | -2.28413 |
| Leu140 | -1.8568 | -0.42722 | 0.380511 | -1.90351 |
| MET158 | -1.88856 | -0.24301 | 0.297095 | -1.83447 |
| Leu109 | -1.83964 | -0.41766 | 0.56906 | -1.68824 |
| Phe144 | -1.90497 | -0.39548 | 0.656424 | -1.64403 |
| Val306 | -1.14166 | -0.05568 | -0.07883 | -1.27618 |
| Ser191 | -0.96024 | -1.15571 | 0.952606 | -1.16335 |
| Met304 | -1.26012 | 0.053739 | 0.09109 | -1.1153 |

**Table S3.** Primers for qRT-PCR.

| **Primers** | **Sequences (5' to 3')** |
|---|---|
| Nn7OMT-F | CATGAAGGTGAGGTCGGTCT |
| Nn7OMT-R | CCAGATTCAGCTTTGCACCG |
| Actin-F | AGGGAGAAGATGACCCAGATTA |
| Actin-R | GTTGTTCTACCACTGGCGTATAG |

**Table S4.** Primers for site-directed mutagenesis of Nn7OMT.

| Primers | Sequence (5'-3') |
|---|---|
| 308A-F | GCTGGTCACAGCAGGTGGTAAAGAAAGAAGCGAAGAAG |
| 308A-R | CTTTACCACCTGCTGTGACCAGCATAGACATGTCCAGA |
| 154A-F | GCTGGTCACAGCAGGTGGTAAAGAAAGAAGCGAAGAAG |
| 154A-R | TCCCCTCGCTTGCCAATCGATTGATCACCGAGTCTTTGCC |
| 307A-F | GCTGGTCACAGCAGGTGGTAAAGAAAGAAGCGAAGAAG |
| 307A-R | TACCACCGGGTGCGACCAGCATAGACATGTCCAGATTCAGC |
| 140A-F | GGGCGAGGACGCAGAGGAATTATTTGGCAAAGACTCGG |
| 140A-R | ATAATTCCTCTGCGTCCTCGCCCTCGTGACACTTCTCA |
| 158A-F | GAGCGAGGGGGCAACGAATCTAACGAGTTTGATGGCGG |
| 158A-R | TTAGATTCGTTGCCCCCTCGCTCAACAATCGATTGATC |
| 109A-F | CTTCGCTCTCGCAATCTTCTACGAGATGGACGCTTGGC |
| 109A-R | CGTAGAAGATTGCGAGAGCGAAGGATGCGAGGTTCTTC |
| 144A-F | AGAGGAATTAGCAGGCAAAGACTCGGTGATCAATCG |
| 144A-R | AGTCTTTGCCTGCTAATTCCTCTGCGTCCTCGCCCTC |
| 306A-F | GTCTATGCTGGCAACACCCGGTGGTAAAGAAAGAAGCG |
| 306A-R | CACCGGGTGTTGCCAGCATAGACATGTCCAGATTCAGC |
| 191A-F | CGTGGGCGGGGCAACTGGGGTAGCTGCGCGTGCCATCG |
| 191A-R | CTACCCCAGTTGCCCCGCCCACGTCAATTAGGGACCCT |
| 304A-F | GGACATGTCTGCACTGGTCACACCCGGTGGTAAAGAAA |
| 304A-R | GTGTGACCAGTGCAGACATGTCCAGATTCAGCTTTGCA |
